# Emerging invasion risks of non-native urban trees in continental Europe under a changing climate

**DOI:** 10.64898/2026.03.16.712173

**Authors:** Mihaela Britvec, Marina Piria, Ivana Vitasović Kosić, S. Luke Flory, Božena Mitić, Sara Essert, Dario Hruševar, Seokmin Kim, Ivica Ljubičić, Lorenzo Vilizzi

**Affiliations:** University of Zagreb Faculty of Agriculture, Department of Agricultural Botany, Zagreb, Croatia; University of Zagreb Faculty of Agriculture, Department of Fisheries, Apiculture, Wildlife Management and Special Zoology, Zagreb, Croatia; University of Lodz, Faculty of Biology and Environmental Protection, Department of Ecology and Vertebrate Zoology, Lodz, Poland; Agronomy Department, University of Florida, Gainesville, USA; Invasion Science Institute, University of Florida, Gainesville, USA; University of Zagreb Faculty of Science, Department of Biology, Division of Botany, Zagreb, Croatia

**Keywords:** climate change, invasive plants, urban trees, Terrestrial Plant Species Invasiveness Screening Kit (TPS-ISK), urban green areas

## Abstract

Urban green areas often harbour numerous non-native urban trees, many of which have characteristics that predispose them to escape from cultivation and become potentially invasive. Climate change is expected to exacerbate this risk by creating favourable conditions for species that are currently climatically restricted. The potential risks for invasiveness of urban tree species in continental Europe are not yet known. Here, we provide a comprehensive risk screening of 34 non-native urban tree species in continental Europe, for both current and projected future climate scenarios. Using the Terrestrial Plant Species Invasiveness Screening Kit (TPS-ISK v2.4), we assessed invasion risk based on biogeography, ecology, and projected responses to climate change. Results showed that under current conditions, 10 species (29.4%) were categorised as high risk, 23 (67.6%) as medium risk and one (2.9%) as low risk. The inclusion of climate change projections increased the number of high risk species to 11, with seven species categorised as very high risk. These taxa exhibit strong ecological plasticity, high reproductive performance and broad environmental tolerance, which together with projected warming, emphasises their significant potential for further spread. Our results emphasise the urgent need for early detection, continuous monitoring and proactive management of non-native urban trees in Europe, especially those that are widely used in horticulture and forestry. By integrating invasion biology with climate change risk screening, this study provides an important basis for evidence-based policy and management strategies to mitigate future ecological and economic impacts of invasions by urban trees.

## Introduction

Today, more than half of the world’s population – over 4 billion people – lives in urban areas (cities, towns and suburbs), a proportion that is expected to increase to 68% by 2050 (UN 2019). Parks and other urban green areas provide aesthetic and cultural services to people and have positive effects on human health and well-being (Chang et al. 2017; WHO 2023). Urban green areas also play an important role in providing important environmental services (e.g. by reducing the impact of urban heat islands and improving air quality) and ecological services (e.g. by providing habitats for urban wildlife and biodiversity conservation) (Paudel and States 2023). However, planted ornamental flora in urban green areas frequently contains a large number of cultivated non-native species (Mayer et al. 2017), which may escape cultivation and invade surrounding natural areas.

Ornamental horticulture is the most important pathway for non-native plant introductions worldwide (Dehnen-Schmutz et al. 2007; Dehnen-Schmutz 2011; Hinsley et al. 2024; Humair et al. 2014). While most of these plants can only survive where planted under intensive management practices (e.g. watering, nutrition), others escape from cultivation (Petřík et al. 2019). After a potential escape, they face the barriers of the naturalization and invasion processes (Richardson and Pyšek 2012), while functional traits of these species (Dyderski and Jagodziński 2019; Williams et al. 2015) as well as climatic and environmental factors act synergistically to influence their success and ability to disperse (Sülle et al. 2023). Among functional traits, a long flowering period, large height, seed mass, total biomass, high germination rate and dispersal ability as well as high stress tolerance enhance the invasion potential of plant species (Richardson and Pyšek 2012). These characteristics are also typical for the species used in horticulture because they facilitate plant establishment and growth in gardens (Sülle et al. 2023). Given increasing urbanization and the constant rise of horticultural practices that favour managed parks and residential gardens, the risk of ornamental plant escape to natural habitats is increasing (Marco et al. 2010).

Since dozens of ornamental plants in temperate regions come from warmer areas (Zieritz et al. 2016), these species often survive and grow in horticultural environments but do not establish self-sustaining populations in the wild. In other words, these non-native ornamental plants are currently still outside their realized climatic niches but are inside their tolerance climatic niches (Sülle et al. 2023). Thus, under climate change (IPCC 2023), they may encounter more suitable environmental conditions on a larger scale, increasing their chance of escaping cultivation and becoming naturalized (Haeuser et al. 2018), and possibly invasive (Appalasamy et al. 2020; Bayón et al. 2022; Bradley et al. 2010; Dullinger et al. 2017; Gaertner et al. 2017; Haeuser et al. 2018; Lososová et al. 2018; Pergl et al. 2016; Petřík et al. 2019; Potgieter et al. 2018).

Early identification of non-native species with invasive potential is critical, as even a single invasive species can significantly disrupt biodiversity and cause substantial economic losses (Hulme et al. 2009). Within risk analysis frameworks, the first step—risk identification—focuses on determining which species are most likely to become invasive in a given region (Vilizzi et al. 2022a). To support this process, electronic decision-support tools have been developed, among which the Aquatic Species Invasiveness Screening Kit (AS-ISK) has become widely applied (Copp et al. 2016, 2021; Vilizzi et al. 2021). Building on this approach, AS-ISK was adapted into the Terrestrial Plant Species Invasiveness Screening Kit (TPS-ISK: Vilizzi et al. 2024), which currently represents the state-of-the-art tool for global application in assessing plant invasiveness.

Although non-native plants have been studied in some urban areas (Alessandrini et al. 2025; Appalasamy et al. 2020; Lososová et al. 2012; Lubarda et al. 2024; Pergl et al. 2016; Petřík et al. 2019; Rat et al. 2017; Ricotta et al. 2012; Wirth et al. 2022), understanding risks of invasiveness of urban trees species in continental Europe is lacking. This research gap is crucial given the challenge posed by climate change, which may increase the invasiveness potential of certain ornamental species, thereby posing an increasing threat to biodiversity (Brandu et al. 2020).

In this study, we focus on urban tree species because they are among the main plant invaders worldwide (Çoban et al. 2021; Čeplová et al. 2017; Richardson and Rejmánek 2011) and some of them are considered as the most harmful invasive species in urban systems (Kowarik et al. 2013; Potgieter et al. 2018). Our goal is to provide the first detailed information on invasion risk of non-native urban trees species in continental Europe under current and future climate conditions. We hypothesize that changing climate would increase the invasion risk of species. In this way, we aim to provide insights into the drivers of non-native species invasions in continental Europe, thereby informing management strategies and contributing to the protection of local ecosystems while mitigating potential economic losses.

## Materials and methods

### Species selection

The list of urban tree species was compiled based on three criteria: (1) an inventory of species from ten parks in Zagreb, Croatia, conducted through field surveys in 2023 and 2024 by the first author, supplemented with data from the Zrinjevac Subsidiary of the Zagreb Holding database; (2) inclusion of species listed in the Royal Horticultural Society Encyclopedia of Plants and Flowers, which provides for a comprehensive reference of plants commonly cultivated in European urban green areas and gardens (Brickell 2019); and (3) selection of species that are non-native to continental Europe, as verified using the Euro+Med Plantbase (https://europlusmed.org/) and the Global Biodiversity Information Facility (GBIF: https://www.gbif.org/) databases. Species were included if they met all three criteria, namely occurrence in urban park inventories, documented use in European urban horticulture, and non-native status in continental Europe.

In total, 34 taxa (33 species and one subspecies) of urban trees were selected, representing some of the more commonly planted trees in European cities (Ossola et al., 2020), and were therefore considered representative for a continental-scale risk screening (Table 1). Taxomomic nomenclature follows the Catalogue of Life Annual Checklist 2025 (https://www.catalogueoflife.org/), the International Plant Names Index (http://www.ipni.org), and Plants of the World Online (https://powo.science.kew.org/). All taxa are hereafter referred to as species.

**Table 1.**
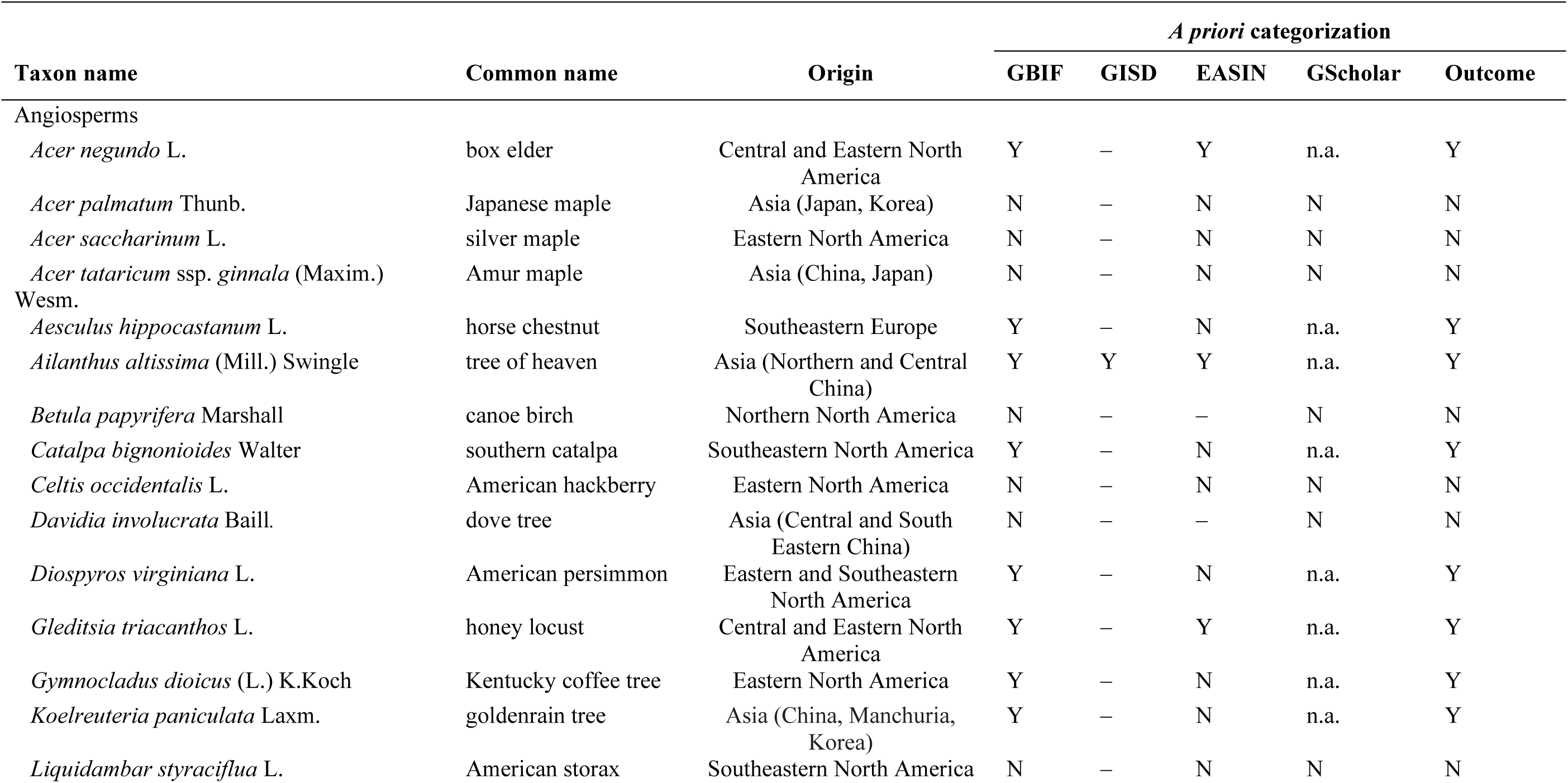
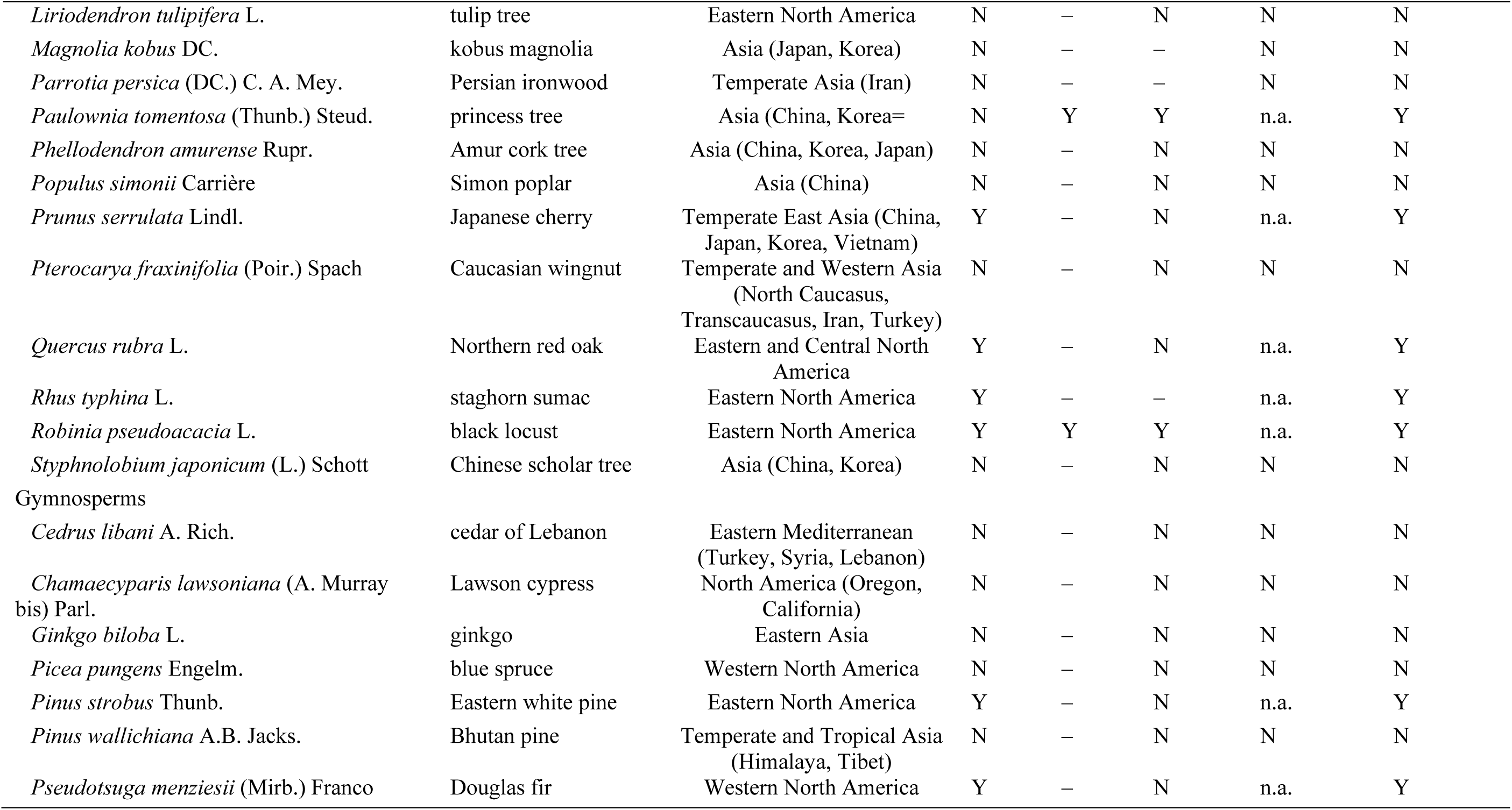
Non-native urban trees (origin: WFO, 2025) screened using the Terrestrial Plant Species Invasiveness Screening Kit (TPS-ISK) for their invasion risk in Continental Europe. *A priori* categorization (Outcome: N = non-invasive; Y = invasive) follows the four-step protocol of Vilizzi et al. (2022b): (1) Global Biodiversity Information Facility (https://www.gbif.org/); (2) Global Invasive Species Database (GISD: www.iucngisd.org); (3) European Alien Species Information Network (EASIN: https://easin.jrc.ec.europa.eu/easin); (4) Google Scholar literature search. N = no impact/threat; Y = impact/threat; ‘–’ = absent; n.a. = not applicable.

### Risk screening

Risk screening was undertaken with the TPS-ISK v2.4.1 (download at https://tinyurl.com/ISK-toolkits). This taxon-generic decision support tool complies with the ‘minimum standards’ for screening non-native species under EC Regulation No. 1143/2014 on the prevention and management of the introduction and spread of invasive alien species. The TPS-ISK consists of 55 questions of which 49 comprise the Basic Risk Assessment (BRA) and six the Climate Change Assessment (CCA). The BRA addresses the biogeography/invasion history and biology/ecology of the species, and the CCA requires the assessor to predict how future predicted climatic conditions are likely to affect the BRA with respect to risks of introduction, establishment, dispersal, and impact. The CCA component is considered an ongoing and cumulative process influencing species performance and spread across Europe.

Screenings were conducted separately by eight assessors (BM, DH, IL, IVK, MB, SE, SK, SLF), with each assessor screening from two to nine species. The completed screenings were then subjected to quality control by MP and LV. All assessors are specialists in the biology and ecology of the urban trees under study. In this respect, ISK screenings do not represent unstructured expert opinion, but a standardized expert elicitation process governed by fixed questions, predefined scoring ranges, confidence rankings, and statistically calibrated risk thresholds. The tool constrains assessor input within a reproducible decision framework, thereby minimizing individual bias. When applied by trained assessors with taxon-specific expertise, this approach has repeatedly demonstrated high internal consistency and transferability across regions and taxa (Vilizzi et al., 2022b). Accordingly, the use of a single trained assessor per species represents an accepted and validated application of the methodology. Previous applications of the ISK family of toolkits have demonstrated high internal consistency across assessors when applied by trained experts, supporting the robustness of single-assessor screenings (Vilizzi et al. 2022b).

Following the protocol by Vilizzi et al. (2022b), the assessor provided for each question a response, a confidence level, and a justification (Vilizzi and Piria 2022). Upon completion of a screening, two outcome scores are obtained: BRA and BRA+CCA. Scores < 1 rank the species as ‘low risk’, whereas scores ≥ 1 rank the species as ‘medium risk’ or ‘high risk’. The distinction between medium and high risk is based on a calibrated threshold obtained by Receiver Operating Characteristic (ROC) curve analysis (Vilizzi et al. 2022a, b). This approach is subject to the availability of a representative number of species (at least 15–20) categorised *a priori* as either invasive or non-invasive in a relatively balanced proportion (Vilizzi et al. 2022b). Given that this criterion was satisfied for the angiosperms but not for the gymnosperms (Table 1), an overall threshold was computed for all urban trees and a separate one for the angiosperms. Additionally, an *ad hoc* threshold (as per Britton et al. 2011) was used to distinguish ‘very high risk’ species within those ranked as high risk.

Fitting of the ROC curves was with pROC (Robin et al. 2011) for R x64 v4.4.0 (R Development Core Team 2024). Permutational ANOVA with normalisation of the data was used to test for differences in the confidence factor (see Vilizzi et al. 2022b) between the BRA and BRA+CCA; this used a Bray-Curtis dissimilarity measure, 9999 unrestricted permutations of the raw data, and with statistical effects evaluated at α = 0.05. Following identification of the threshold score, evaluation of the risk classifications to identify false-positive and false-negative rankings was not applied to the medium-risk species because their further evaluation in a follow-up risk assessment depends on management priorities and the financial availability. All screenings were carried out under the supervision of LV, who was responsible for quality control of the biological/ecological data and of the methodological (i.e. database-related) aspects of the study.

## Results

For all tree species, the ROC curve resulted in an AUC of 0.6947 (0.5019–0.8876 95% CI) and a threshold of 41. For the angiosperms, the ROC curve resulted in an AUC of 0.5893 (0.3548–0.8238 95% CI) and again in a threshold of 41. The AUC values were overall within the acceptable limits for the discriminatory power of the test (Hosmer et al. 2013). This threshold was therefore used for calibration of the risk outcomes to distinguish between medium-risk and high-risk species (Table 2; refer to the Supplementary online material Appendix S1 for reports of the 34 screened species).

**Table 2.**
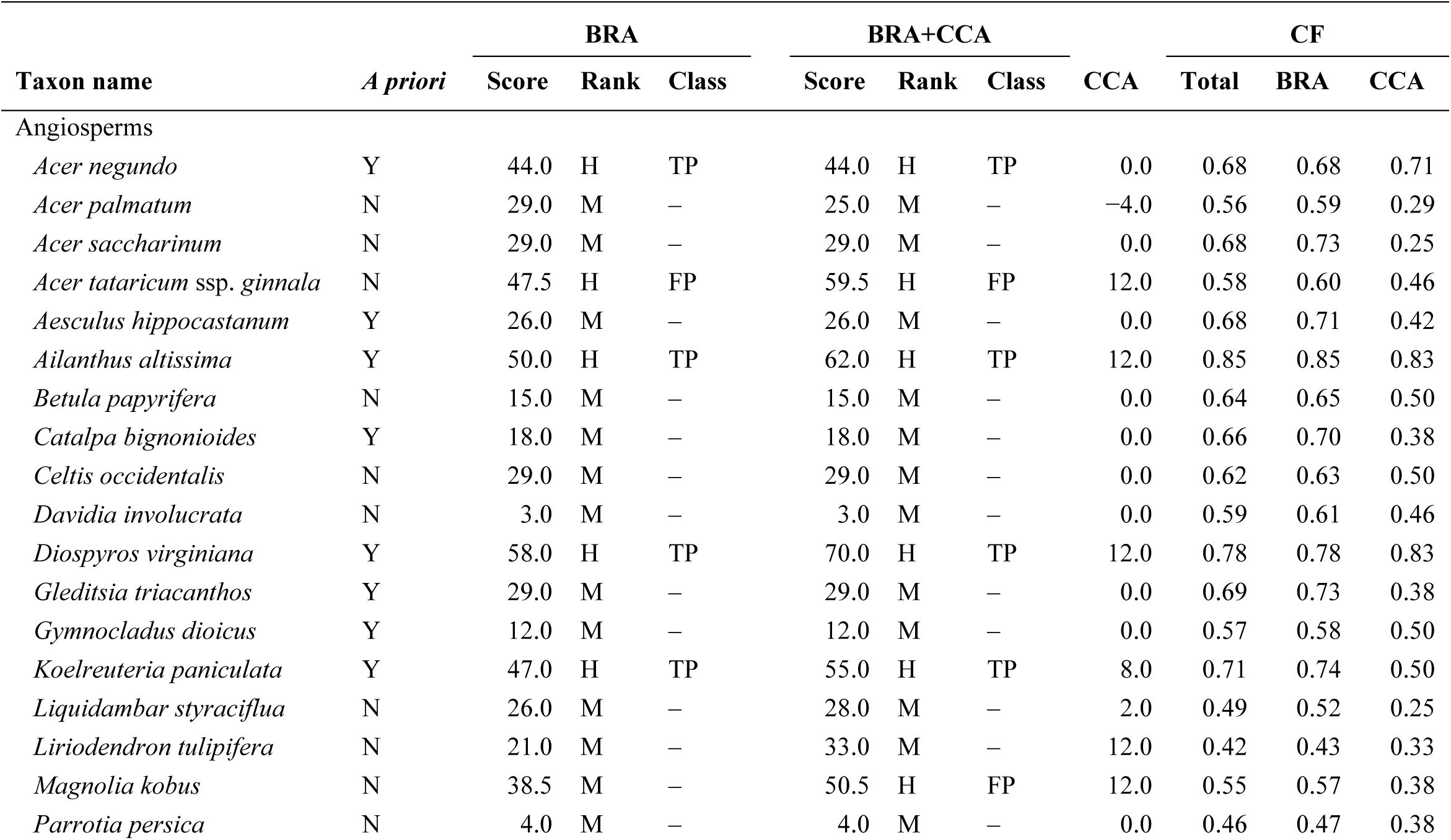
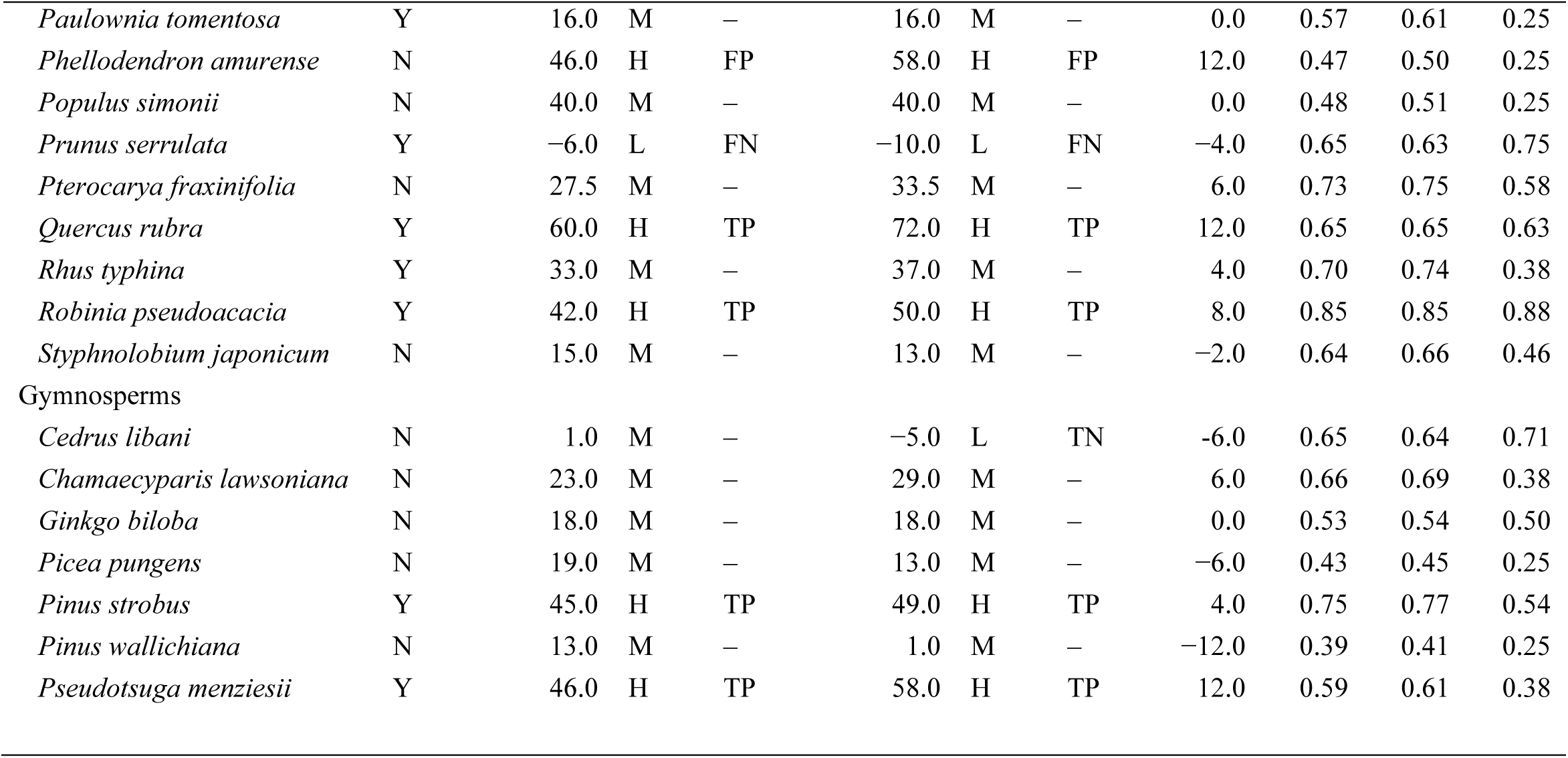
Risk outcomes for the non-native urban trees screened with the TPS-ISK for Continental Europe. For each taxon, the following information is provided: *a priori* categorisation of invasiveness (N = non-invasive; Y = invasive: see Table 1), Basic Risk Assessment (BRA) and BRA + Climate Change Assessment (BRA+CCA) scores with corresponding risk ranks based on a calibrated threshold of 41 (L = Low; M = Medium; H = High; VH = Very high, based on an *ad hoc* threshold ≥ 50: see text for details), classification (Class: FP = false positive; TN = true negative; TP = true positive; ‘–’ = not applicable as medium-risk: see text for details), CCA as difference between BRA+CCA and BRA scores, and confidence factor (CF). Risk outcomes for the BRA scores (within interval): L [−20, 1[; M, [1, 41[; H]41, 50[; VH [50, 72]. Risk outcomes for the BRA+CCA scores: L, [−32, 1[; M [1, 41[; H]41, 50[; VH [50, 82]. Note the reverse bracket notation indicating an open interval.

Based on the BRA scores (Table 2, Fig. 1a), 10 (29.4%) species were ranked as high risk, 23 (67.6%) as medium risk and 1 (2.9%) as low risk. Of the 15 species categorised *a priori* as invasive, eight were ranked as high risk (true positives: box elder *Acer negundo*, tree of heaven *Ailanthus altissima*, American persimmon *Diospyros virginiana*, goldenrain tree *Koelreuteria paniculata*, Eastern white pine *Pinus strobus*, Douglas fir *Pseudotsuga menziesii*, Northern red oak *Quercus rubra*, black locust *Robinia pseudoacacia*) and one as low risk (false negative: Japanese cherry *Prunus serrulata*). Of the 19 species categorised *a priori* as non-invasive, two were ranked as high risk (false positives: Amur Maple *Acer tataricum* ssp. *ginnala,* Amur cork tree *Phellodendron amurense*). Of the 23 medium-risk species, 17 were *a priori* non-invasive and six invasive.

**Figure 1.**
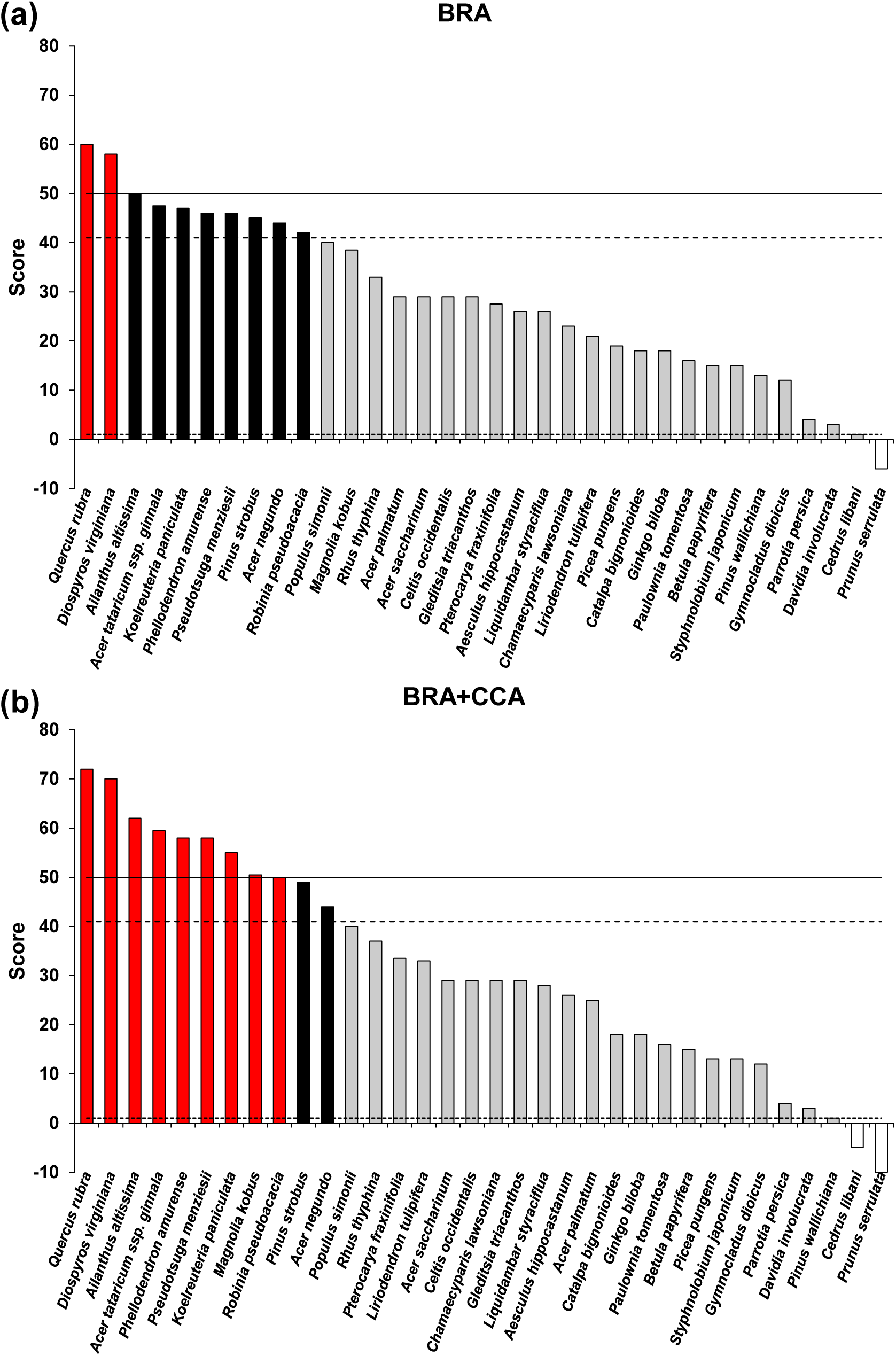
Risk outcome scores for the non-native urban trees screened with the Terrestrial Plant Species Invasiveness Screening Kit (TPS-ISK) for Continental Europe. (a) Basic Risk Assessment (BRA) scores; (b) BRA + Climate Change Assessment (BRA+CCA) scores. Red bars = very high-risk species; Black bars = high-risk species; Grey bars = medium-risk species; White bars = low-risk species; Solid line = very high-risk threshold; Hatched line = high-risk threshold; Dotted line = medium-risk threshold. Thresholds as per Table 2.

Based on the BRA+CCA scores (Table 2, Fig. 1b), 11 (23.5%) species were ranked as high risk, 21 (17.6%) as medium risk and 2 (2.9%) as low risk. Ranking of the *a priori* invasive species was the same as per the BRA. Of the *a priori* non-invasive species, three were ranked as high risk (same species as per BRA plus Kobus magnolia *Magnolia kobus*) and one as low risk (same species as per BRA). Of the 21 medium-risk species, 15 were *a priori* non-invasive and six invasive.

Based on an *ad hoc* threshold ≥ 50, there were two very high-risk species for the BRA and BRA+CCA (*Diospyros virginiana* and *Quercus rubra*), and seven for the BRA+CCA only (Amur maple *Acer tataricum* ssp. *ginnala*, *Ailanthus altissima*, goldenrain tree *Koelreuteria paniculata*, *Magnolia kobus*, *Phellodendron amurense*, Douglas fir *Pseudotsuga menziesii*, *Robinia pseudoacacia*) (Table 2, Fig. 1a, b). The CCA resulted in an increase in the BRA score (cf. BRA+CCA score) for 15 (44.1%) species, in no change for 13 (38.2%) and in a decrease for 6 (17.6%) (Table 2).

The mean CF_Total_ was 0.616 ± 0.020 SE, the mean CF_BRA_ 0.635 ± 0.020 SE and the mean CF_CCA_ 0.463 ± 0.032 SE. The mean CF_BRA_ was higher than the mean CF_CCA_ (*F*^#^ = 0.94, *P^#^* = 0.333; # = permutational value).

## Discussion

### Risk outcomes

Overall, the present results indicate that the risk screening satisfactorily differentiated species by invasion potential, with most species classified as medium risk and about one-quarter as high risk. The model’s performance metrics suggest moderate predictive accuracy for distinguishing between medium- and high-risk species. The BRA and BRA+CCA models consistently ranked known invasive species such as *Ailanthus altissima*, *Diospyros virginiana*, and *Quercus rubra* among the highest risk categories. The CCA component slightly increased risk scores for several species, emphasizing how future climatic conditions may heighten invasion risks.

*Ailanthus altissima*, native to northeast and central China and Taiwan, represents one of the most widespread invasive tree species globally (Kowarik and Säumel 2007). First introduced to Europe in the 18th century, it has since spread rapidly due to its resilience to herbivory, tolerance of urban conditions, and popularity as an urban and avenue tree (Swingle 1916; Kowarik and Böcker 1984; Kowarik and Säumel 2007; Zaraś-Januszkiewicz et al. 2014). Today, *A. altissima* colonizes both heavily disturbed urban environments and semi-natural habitats, including abandoned industrial sites, roadsides, riverbanks, forest edges, and agricultural margins (Karlović and Prebeg 2020). Its invasive success is attributed to prolific seed production (Kowarik and Säumel 2007; Soler and Izquierdo 2024), root suckering (Constán-Nava et al. 2010), rapid growth, early maturity (Terzi et al. 2021), broad environmental tolerance (Sladonja et al. 2015), allelopathic effects (Heisey 1996), and release from natural enemies (Meloche and Murphy 2006). These traits, combined with anthropogenic disturbance and climate change, strongly promote persistence and spread (Isler et al. 2023). While temperature thresholds constrain establishment in cooler regions and at higher altitudes (> 900 m a.s.l.) (Motti et al. 2021), the climatic overlap between its native range and southern Europe suggests continued expansion (Zaraś-Januszkiewicz et al. 2014).

*Diospyros virginiana*, native to the eastern and southeastern United States (POWO 2025), has so far shown limited spread in Europe, where it was introduced in the late 19th and early 20th centuries for ornamental and fruit purposes (Briand 2005; Kluge and Tessmer 2018; Burge 2018). In Croatia, its presence dates to 1859, when it was planted in the Trsteno Arboretum (Idžojtić et al. 2013). While natural regeneration is mostly restricted to abandoned and ruderal sites (Marić et al. 2024), its traits—high seed viability, endozoochoric dispersal by birds and mammals, tolerance of drought and poor soils, and deep taproot systems (Dirr 2009)—indicate resilience. Climate change may amplify its potential by reducing frost damage and enabling establishment in formerly unsuitable regions (Walther et al. 2009). Although its invasive behaviour remains localized, projections suggest that warmer conditions, as present in Mediterranean region, may elevate its risk from locally naturalized to regionally established (Marić et al. 2024).

*Quercus rubra* was introduced from North America in 1691 and is now widespread across Europe, covering more than 350,000 ha, particularly in France, Germany, Poland, Hungary, and Slovenia (Nicolescu et al. 2020b; Májeková et al. 2023). Initially valued as an urban tree, its use later expanded to forestry and reforestation due to its rapid growth, high-quality timber, and ecological flexibility (Dyderski et al. 2020). Compared with native oaks (*Quercus robur* and *Q. petraea*), *Quercus rubra* exhibits higher tolerance to drought and other environmental stresses, enhancing its competitiveness under climate change (Bauweraerts et al. 2014; Nicolescu et al. 2020b). Its regeneration relies both on acorn dispersal by animals (up to 1.5 km from parent trees) and strong sprouting ability (Dyderski et al. 2020; Nicolescu et al. 2020b). Given these advantages, *Quercus rubra* is projected to increasingly replace declining native oak species in forests in the future (Dyderski et al. 2020).

Other species scored as high invasion risk, including *Acer negundo, A*. *tataricum* ssp*. ginnala, Koelreuteria paniculata, Phellodendron amurense, Pinus strobus, Pseudotsuga menziesii,* and *Robinia pseudoacacia*, were also found to pose significant risks under both present and future climate conditions. *Acer negundo* and *A. tataricum* ssp*. ginnala* spread via wind- and water-dispersed samaras, thrive on a wide range of soils, and display strong drought tolerance, making them successful colonizers of urban and disturbed habitats (Dirr 1997; Porté et al. 2011; Kostina et al. 2014; Sikorska et al. 2019; Szabó et al. 2022; Ovcharova et al. 2024). *Koelreuteria paniculata* and *Phellodendron amurense* are at earlier stages of naturalization, but their reproductive strategies and climate adaptability highlight future risks (Idžojtić 2013; Zhao et al. 2023; O’Connell et al. 2023). *Pseudotsuga menziesii*, while valued in forestry, can regenerate naturally and alter soil biota (Schmid et al. 2014; Bayón et al. 2022). *Pinus strobus* and *Robinia pseudoacacia* are already invasive in several European countries, with the latter exerting particularly strong ecosystem impacts through nitrogen fixation, allelopathy, and competitive stand formation (Mandak et al. 2013; Vitkova et al. 2017; Nicolescu et al. 2020a).

Overall, these findings indicate that a combination of ecological traits, reproductive strategies, and increasing climatic suitability drives the invasion potential of both currently widespread and still-emerging species. While *Ailanthus altissima, Diospyros virginiana*, and *Quercus rubra* present the highest immediate risks, several other taxa require close monitoring to prevent future large-scale impacts on European biodiversity.

### Climate change

Climate change is expected to play a central role in shaping the invasive potential of non-native tree species in Europe. Climate change is often discussed in the context of rising mean temperatures; however, it encompasses a broader set of environmental changes, including shifts in precipitation regimes, increased frequency and intensity of extreme events, altered seasonality, and changes in climatic variability (van der Wiel and Bintanja 2021; Wood 2023). These multiple dimensions can interact with land use, disturbance, and biotic interactions to influence species performance and distribution in complex and sometimes non-linear ways (Fernández de Castro et al. 2018). In this study, climate-change effects are incorporated in a simplified manner that aligns with the screening-level objective of the method. The approach used in this study provides an overall estimate of invasion risk rather than a process-based representation of individual climatic drivers. Consequently, it does not disentangle the roles of temperature, precipitation, green house gas emissions or their interactions, and the results of this study should be interpreted as a first-pass assessment and complemented by spatially explicit analyses for management purposes.

Climate-change impacts vary substantially across Europe, which is climatically heterogeneous (Dyderski et al. 2018). Species may benefit from changing conditions in some regions, such as *Ailanthus altissima* for southern Europe, while may experience reduced suitability and range contraction elsewhere (Isler et al. 2023). The modelling studies for *Ailanthus altissima*, suggest that warmer and drier conditions will support its persistence and expansion, particularly in drought-prone habitats (Isler et al. 2023). Although its spread is still limited by temperature at higher altitudes (>900 m a.s.l.) and in colder regions (Motti et al. 2021), the overlap between climatic conditions in its native range and those of southern Europe indicates considerable scope for further range expansion (Zaraś-Januszkiewicz et al. 2014).

*Diospyros virginiana* may also benefit from ongoing climate change. Rising winter temperatures and longer growing seasons could reduce frost damage, improve seedling survival, and open up regions that were previously unsuitable for its establishment (Walther et al. 2009; Marić et al. 2024). Its inherent drought tolerance and capacity to persist on nutrient-poor soils further increase its resilience under future climate stress. For *Quercus rubra*, warming scenarios may strengthen its competitive advantage over native European oaks. Experimental studies indicate that its greater drought tolerance compared with *Quercus robur* and *Q. petraea* could allow it to replace declining native populations in future European forests (Bauweraerts et al. 2014; Dyderski et al. 2020).

These dynamics are not limited to the three the highest scored species. Climate change is also predicted to enhance the establishment of other non-native trees currently present in Europe. For example, warmer conditions and longer growing seasons may increase seed viability and colonization success in *Koelreuteria paniculata* (Ljubojević et al. 2021), while facilitating the spread of *Phellodendron amurense* beyond its native climatic range (Zhao et al. 2023). Also, future climatic conditions will likely favor the occurrence of *Robinia pseudoacacia* in Central and Northeastern Europe where this species is still absent or relatively rare (Puchałka et al. 2021). Similarly, both *Pseudotsuga menziesii* and *R. pseudoacacia* are projected to benefit from increased drought tolerance, which could enhance their competitiveness in European forests under future conditions (Dyderski et al. 2018; Lange et al. 2022).

Urban centers are consistently warmer and often have longer growing seasons than nearby rural areas, which alters species phenology and thermal regimes relevant to establishment (Abulibdeh 2021; Carlon et al. 2024). In European cities, urban heat island intensities can exceed ∼8 °C under certain conditions, and urbanization has been shown to extend growing seasons relative to surrounding rural areas, with climatic influences detectable well beyond the urban core along urban–rural gradients (Dallimer et al. 2016; Mentaschi et al. 2022). Such conditions may facilitate the establishment of cultivated ornamental species, including urban trees, by reducing climatic constraints, particularly cold limitation, and by exposing populations to warmer and more variable conditions than those in surrounding landscapes (Ariori et al. 2017). As a result, urban environments may act as initial establishment foci and potential pre-adaptation zones, enabling species to persist beyond cultivation and increasing the likelihood of subsequent spread as regional climates warm (Carlon et al. 2024). Although these urban-specific effects are not explicitly captured in our analysis, they highlight an additional pathway through which climate change and land use may interact to influence invasion risk.

Overall, climate change, particularly in urban centres, is likely to increase colonization opportunities for a wide range of non-native tree species. While it may amplify the already substantial invasive potential of *Ailanthus altissima* and *Quercus rubra*, it could also elevate species such as *Diospyros virginiana*, *Koelreuteria paniculata*, and *Phellodendron amurense* from locally naturalized or regionally constrained taxa to more widespread and ecologically significant invaders.

### Management considerations

Management of non-native tree species in Europe remains highly context-dependent, reflecting both ecological risks and socio-economic values. The EU List of Invasive Alien Species of Union concern, established under Regulation (EU) 1143/2014 and its subsequent amendments, represents the primary legislative framework for coordinated management. However, only a limited number of tree taxa are currently included. For example, *Ailanthus altissima* was added in 2019 under Commission Implementing Regulation (EU) 2019/1262 (EU 2022), following widespread evidence of its aggressive spread and ecological impacts.

By contrast, several other non-native tree species with documented invasive potential remain excluded from the Union list. *Quercus rubra*, which is considered invasive or potentially invasive by some authors and official bodies (Chmura 2020), is not included in the Union list despite widespread plantings across Europe (European Union Regulation 2022/1203). *Acer tataricum* ssp. *ginnala* illustrates a comparable case: while it is an emerging invasive threat in North America (Schuster and Reich 2018) and has been classified as high-risk in Latvia (Evarte-Bundere et al. 2022), it has not been recognized as a species of Union concern (EU 2022). *Diospyros virginiana* presents a different challenge. Its spread is still limited, but local escapes, for example in Croatia (Marić et al. 2024), demonstrate the importance of early detection and monitoring. Restricting its further introduction into vulnerable ecosystems and promoting awareness among horticultural stakeholders may prevent future invasiveness.

The same uncertainty applies to species such as *Koelreuteria paniculata*, *Phellodendron amurense*, *Pseudotsuga menziesii*, *Pinus strobus*, and *Robinia pseudoacacia*. Although they are not currently listed at the Union level, evidence of local naturalization and ecological impacts suggests the need for proactive monitoring and risk assessment (Boršić et al. 2016; Nicolescu et al. 2020a).

In conclusion, these examples highlight the need for continuous monitoring and risk assessment, even for species not yet included in the EU List of Invasive Alien Species of Union Concern. Particular attention should be paid to taxa that are still freely available in horticultural trade (*Koelreuteria paniculata*, *Phellodendron amurense*), since urban plantings often act as stepping stones for subsequent escape into semi-natural and protected habitats. Integrating ecological, climatic, and socio-economic perspectives will be essential in determining management priorities and mitigating potential long-term impacts on European biodiversity.

## Acknowledgements

The authors would like to thank Martin Veža, MSc. agr., from the Zrinjevac Subsidiary of the Zagreb Holding for providing data of ornamental plants in parks of Zagreb.

## Author contributions

Study conceptualisation: M. Piria, L. Vilizzi; Data preparation: M. Britvec, S. L. Flory, B. Mitić, S. Essert, D. Hruševar, S. Kim, I. Ljubičić, I. Vitasović Kosić; Data analysis: L. Vilizzi; Writing the manuscript: M. Britvec, M. Piria, L. Vilizzi, S. L. Flory, B. Mitić, S. Essert, D. Hruševar, S. Kim, I. Ljubičić, I. Vitasović Kosić

## Data availability statement

All required data/information are provided in the manuscript.

## Declaration of generative AI and AI-assisted technologies in the manuscript preparation process

During the preparation of this work the authors used ChatGPT only to assist with linguistic revision of the manuscript. All scientific content was generated by the authors, who critically reviewed and take full responsibility for the final version.

## Supplementary online material

TPS-ISK report for the 34 screened taxa.

